# Predictive processing for sensory and action neural representations

**DOI:** 10.1101/2025.03.26.645167

**Authors:** Gabriela Iwama, Assaf Breska, Randolph F. Helfrich

## Abstract

Prior information, such as predictive cues, shape perceptual decisions. Predictive theories make conflicting claims about whether the brain preferentially processes expected, as proposed by Bayesian accounts, or unexpected information, as proposed by cancellation accounts. These conflicting mechanisms may co-exist if they differ across sensory and action systems, or if their influence changes as the decision unfolds. To test these alternatives, we tracked sensory and action representations in EEG during a motion-discrimination task with changing cue validity. Cross-temporal decoding revealed distinct stabilization of sensory and action representations. Under strong priors, stabilization of unexpected actions predicted stronger bias by the cue, consistent with cancellation accounts. Unlike action, stabilization of expected sensory information was both positively and negatively correlated with precision-weighted prior bias, depending on the decision stage (before or after stimulus onset), compatible with both Bayesian and cancellation accounts. Together, these findings suggest that predictive processing does not uniformly favor expected or unexpected information. Instead, its effects depend on representational domain and decision stage.

**Statement of Significance:** Predictions help the brain anticipate what will happen by facilitating expected information. However, the brain must stay sensitivity to unexpected information in case predictions are wrong. We show that these two mechanisms can coexist: predictions stabilized sensory and action representations differently, with opposing behavioral predictions across stages of decision formation. Thus, rather than having a universal effect, predictive processing may rely on complementary mechanisms to support flexible behavior.

## Introduction

Because sensory evidence is often incomplete, adaptive behavior depends critically on the ability to generate and use predictions (Knill & Richards, 1996; Körding & Wolpert, 2004; Summerfield & de Lange, 2014). Predictions can disambiguate the interpretation of noisy sensory input and facilitate action preparation, but they can also introduce biases by overweighting prior expectations. For example, when looking for a friend in a crowd, expecting them to wear a red jacket can help distinguish them from other people (sensory evidence) and prepare to greet them (action). While the behavioral impact of predictions is well-characterized, predictive processing theories make competing predictions about whether expected or unexpected information is preferentially facilitated (de Lange et al., 2018). *Bayesian theories* posit that optimal decision-making should weigh prior predictions and bias choices towards expecting outcomes (Wei & Stocker, 2015), and should enhance the neural representations of expected information (Kok et al 2012; Press et al., 2020). In contrast, *cancellation theories* propose that expected information is suppressed (or cancelled) to prioritize the processing of surprising inputs (Bays & Wolpert, 2007; Blakemore et al., 1998; Press et al., 2020).

Despite these conflicting predictions, these mechanisms may not be mutually exclusive. Two possibilities could reconcile these different accounts. First, predictive mechanisms may differ across sensory and action domains. Sensory processing relies on probabilistic inference about latent states (Friston, 2010; Knill & Richards, 1996; Rao & Ballard, 1999). From this perspective, predictions should facilitate expected sensory representations, for example by sharpening tuning curves or increasing pattern separation (Kok et al., 2012; Nigam & Schwiedrzik, 2024; Park et al., 2023; Summerfield & Koechlin, 2008). In contrast, action processing relies on optimal feedback control as the motor system continuously compares the desired movement trajectory to the actual, unfolding trajectory, and uses the difference between them to adjust the movement (Doya, 2021; Scott, 2004, 2012; Wolpert et al., 2011; Yamauchi et al., 2025). Consistent with this feedback control function, the brain suppresses expected sensory consequences of its own actions (Blakemore et al., 1998; Kilteni et al., 2025; Kilteni & Ehrsson, 2017; Shergill et al., 2013). Because the expected consequences are subtracted out, what remains is the prediction error (the unexpected part of the signal), which can be used to correct the motor command. As a result, Bayesian accounts are more prominent in the perceptual literature, whereas cancellation theories are more prevalent in the action literature. Notably, some studies have provided contradictory results, such as weaker representations of expected visual stimuli (Richter et al., 2018), or enhanced representations of sensory signals when they were congruent with hand movements (Yon et al., 2018). Reconciling Bayesian and cancellation theories requires simultaneously investigating predictive mechanisms across sensory and action domains. Second, Bayesian and cancellation mechanisms may co-exist if they operate at different stages of decision-making. As decisions unfold over time, the decision criterion can shift as more sensory information becomes available. For example, when searching for a friend in a crowd, an early glimpse of a red jacket might bias you to mistake a stranger for your friend, but closer inspection can override that initial belief. Thus, the balance between top-down expectations and bottom-up input must be continuously updated (Glaze et al., 2018; Piet et al., 2018; Ruesseler et al., 2023), especially in volatile environments where the reliability of predictions changes over time. The Opposing Process Theory (OPT, Press et al., 2020) offers a temporal account in which predictions initially facilitate expected information. However, if expectations are violated, then predictions facilitate unexpected information.

To test how predictions shape the temporal evolution of sensory and action representations, participants performed a cued motion-discrimination task in which cue validity changed without explicit warning. Thus, participants had to continuously infer the current cue-outcome contingency and decide whether, and to what extent, to rely on prior information. This design allowed us to vary the reliability of prior information while holding sensory evidence constant, enabling us to compare neural representations when priors were weak or strong, and when they were either confirmed or violated. We hypothesized that the stability of predictive representations depends on prior strength and domain (sensory vs. action). First, we predicted that strong priors stabilize sensory and action information more than weak priors. Second, when priors are strong, we expected different effects across domains: priors should stabilize expected sensory representations (Bayesian accounts), but stabilize unexpected action representations (cancellation accounts). Third, we predicted that these dynamics would relate to individual differences in reliance on priors, such that participants who rely more on priors should show stronger stabilization of expected sensory representations and unexpected action representations.

## Materials and methods

### Participants

20 healthy volunteers (10 females, aged 20-31 years, 19 right-handed) were recruited from the student community. All had normal or corrected vision and no history of photosensitive epilepsy, claustrophobia, color blindness, or neurological disorders. Participants provided informed written consent to participate in the study and were paid for their participation according to their performance. This study was approved by the Ethics Committee of the Medical Faculty Tübingen (protocol number 049/2020B02) and conducted in accordance with the Declaration of Helsinki.

### Experimental Design

Before the main task, motion coherence thresholds for the Random Dot Kinematogram (RDK) were estimated using QuestPlus, an adaptive psychometric method (Watson, 2017; Jones, 2018) to yield 75% accuracy. Motion stimuli consisted of white dots (size: .06 degrees of visual angle; density: 16.7 dots/degree; velocity: .03 degrees of visual angle/second; lifetime: 12 frames) moving within a 2.5° circular aperture centered on a fixation cross against a black background. Most dots moved randomly, while up to 20% coherently moved left or right. Participants sat 120 cm from a 28” LED screen (60 Hz, 2560×1440 px) in a dark, magnetically and acoustically shielded room. Responses were made using two response boxes (one per hand). Stimuli were presented using PsychToolbox for MATLAB (Brainard, 1997; Pelli, 1997).

### Behavioral Task

The task combined reversal learning with a motion direction discrimination paradigm. Each trial required three sequential decisions: cue choice, confidence judgment, and motion discrimination. Trial events were interleaved with fixation crosses (see Fig. 1A) and trial onset was self-paced. Cue identity was color-coded (blue or orange). After choosing a cue, participants rated their confidence (low/high) that the cue would be valid (indicate the correct upcoming motion direction in the current trial). A directional cue (left/right arrow) was then presented for 300 ms. To prevent the formation of directional pre-cue bias, the cue direction was randomized with equal probability (50%), regardless of its validity. Therefore, even when participants formed strong priors about the cue validity, this effectively dissociates prior strength from directional prior before the cue presentation for the current trial. This design prevents the formation of expectations about specific cue identities and directions, e.g., p(left|orange) > p(right|orange), and long-term priors, e.g., p(left) > p(right). Finally, participants reported the perceived motion direction (left/right) by pressing a response button with the corresponding (left/right) thumb.

**Figure 1.**
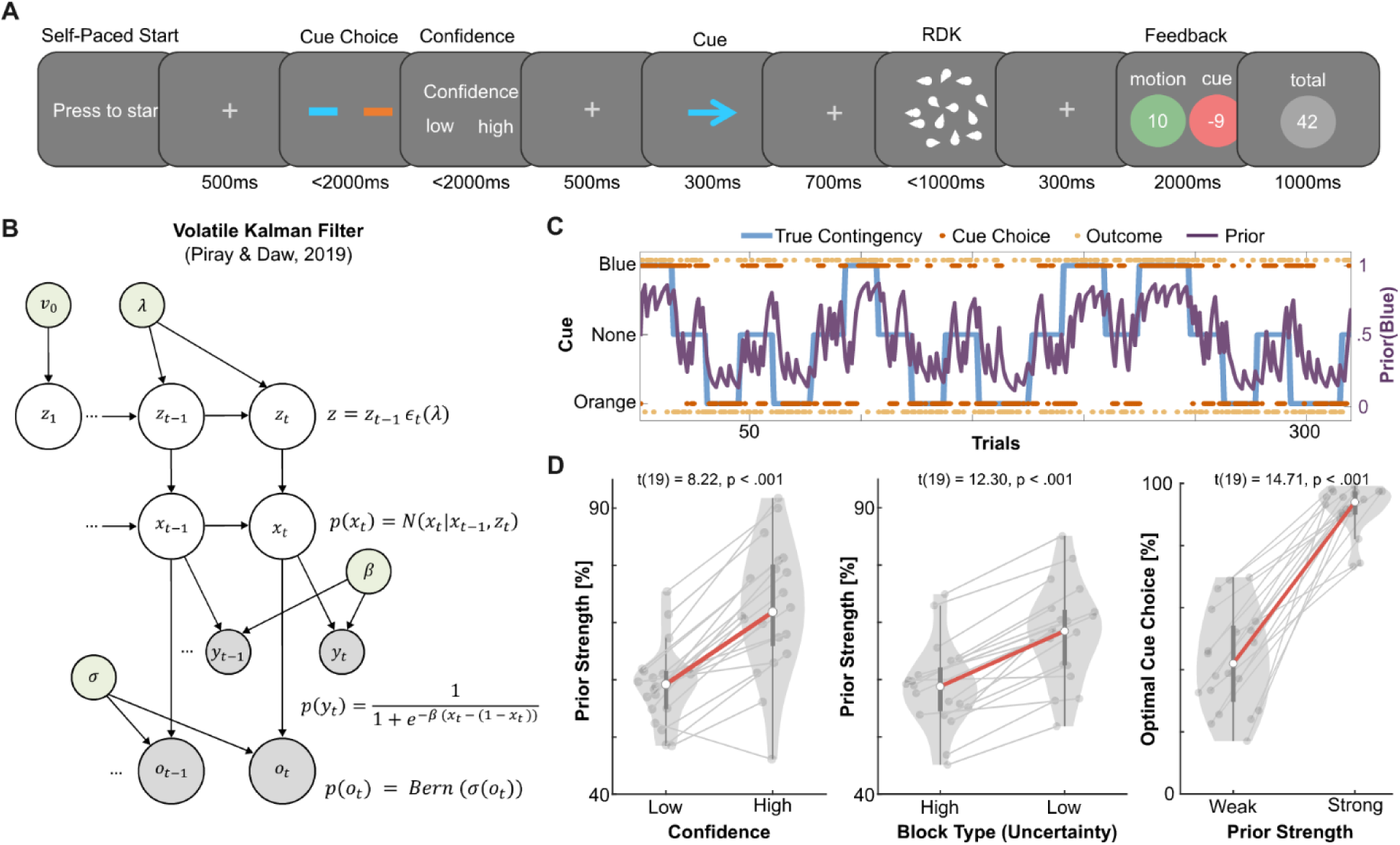
Behavior experiment, modeling, and validation of prior strength. (**A**) Participants performed a cued motion discrimination task with changing cue validities. On every trial, participants selected a cue (orange/blue) and rated their confidence (low/high) that the chosen cue would be valid. The chosen cue randomly indicated left or right. After a delay, participants were presented with the RDK at their individual discrimination level (75% accuracy) and indicated the perceived motion direction (left/right). Feedback was given on both motion accuracy and cue validity. For optimal behavior, participants tracked the validity of the cues across trials. (**B**) We used the Volatile Kalman Filter model (Piray and Daw, 2020) to estimate the evolution of latent priors across trials based on cue choices (orange/blue) and outcome (valid/ invalid). This model consists of two hidden Markov chains: *x_t_* denotes whether the blue cue is valid, and *z_t_* denotes the inverse volatility of *x_t_*. The outcome, *o_t_*, indicates the observed cue validity. The choice, *y_t_*, is given by a softmax function of *x_t_* with a free parameter, *β*, to model how rationally participants follow their prior estimates. Free parameters *λ*, *σ*, and *v*_0_ indicate the (inverse) volatility update rate, observation noise, and initial volatility. (**C**) Single subject example of valid cue (true contingency; blue), cue choice (*y_t_*, orange), outcome (*o_t_*; yellow), and estimated prior (*x_t_*; purple) across trials are shown. Estimated priors closely tracked reversals in the true contingency. (**D**) Estimated prior strength according to confidence (left), block type (middle), and proportion of optimal cue choice during low uncertainty blocks (right). Dots represent individual participants, violin plots show median, interquartile range (IQR), and ±1.5 *IQR*.

Participants were informed that cue validities were probabilistic and could change over time, but not how often these changes would occur. Participants were instructed they could freely decide when to rely on the cue information (i.e., the prior) to guide their motion discrimination judgments. Trials were covertly organized into blocks of low and high uncertainty. During low uncertainty blocks, one of the two cues had 80% probability to correctly indicate the upcoming motion direction (high validity), while the other had 30% probability (low validity). To prevent the formation of long-term cue choice bias, e.g., p(orange) > p(blue), the identity of the high validity cue was pseudo-randomized such that it was never the same for more than two consecutive low uncertainty blocks. To induce states of high uncertainty, low uncertainty blocks were interleaved with blocks in which both cues had low validity (high uncertainty blocks; p(orange) = p(blue). Cue validity on individual trials was pseudo-randomized such that the exponential moving average over the previous 5 trials remained within ±15% of the true cue validity. Low uncertainty blocks lasted 15–30 trials. After 15 trials, transitions to a high uncertainty block occurred if the learning criterion was reached (if the exponential moving average of the cue accuracy over the past 10 trials exceeded 80%). Participants completed 15 training trials in which no validity reversals occurred to familiarize themselves with the trial structure and timing. The main task consisted of up to 320 trials.

We adopted an 80-30% validity structure rather than the classical complementary structure (e.g., 70-30%) used in reversal learning paradigms (Bartolo & Averbeck, 2020; Cools et al., 2002; Wang et al., 2025), because complementary contingencies would lead to equally informative cues. However, even in an 80-30% validity structure, the low validity cue remains (negatively) informative. To discourage the use of information contained in low validity cues, we designed a dual reward system (see “Reward Structure” section), providing separate rewards for accurate motion discrimination and cue validity in that trial (regardless of whether it is the true high valid cue), which promotes choosing high valid cues based on learning while reduces the utility of exploiting anti-correlations. Furthermore, because the true cue validity and reversal volatility were both hidden, this reward structure made it unlikely that participants could reliably learn and use this anti-correlation to predict the upcoming motion direction. If participants had done so, their motion discrimination accuracy would have been higher during high uncertainty blocks than when they used sensory information alone. Consistent with the idea that participants were unable to use the anti-correlation, their motion discrimination accuracy during high uncertainty blocks was not above the calibrated 75% discrimination threshold (p = .33), indicating that participants did not use the anti-correlation information to guide their motion discrimination choices even when both cues had low validities.

### Reward Structure

To balance motion accuracy and cue validity learning, we used a dual-reward system. Participants viewed a feedback screen at the end of the trial, which provided two rewards: one for the motion and another for the cue choice. Then points were awarded for correct motion responses, whereas 10 points were deducted for incorrect responses. Cue choice rewards were scaled by their confidence ratings: high confidence yielded a high-risk, high-reward state, while low confidence entailed safer but smaller rewards. This structure was based on a proper scoring rule (Carvalho et al., 2018), designed to discourage indiscriminate high-confidence ratings. If the cue was valid, participants received 10 points if their confidence was high or 8 points if their confidence was low. For invalid cues, participants lost 9 points if their confidence was high or 4 points if their confidence was low. This design incentivized participants to report low confidence when they were uncertain about which cue had the highest validity. To promote the reliance on prior information, participants were penalized (−20 points) if they did not respond within 1000 ms.

### Behavioral Modeling

To model latent prior strength across trials, we used the Volatile Kalman Filter (VKF; Piray and Daw, 2020), a variant of the classical Kalman filter (Kalman, 1960). Its generative model assumes that the two hidden Markov chains, in which the latent prior indicates the optimal cue for a given trial, *x_t_*, diffuses over time according to the hidden (inverse) volatility of the environment, *z_t_*, such that *x_t_* = *N*(*x_t_*|*x_t_*_−1_, *z*^−1^). In turn, *z_t_* is given by its previous values and evolves dynamically according to a multiplicative diffusion noise, *z_t_* = *z_t_*_−1_*ε_t_*(*λ*). *ε_t_* is defined as a rescaled beta distribution such that *ε_t_* ∈ [0, (1 − *λ*)^−1^], and 0 < *λ* < 1. Therefore, the free parameter *λ* controls the diffusion of the volatility. On the first trial, the inverse mean of *z_t_* is given by a free parameter *v*_0_. The choice, *y_t_*, is defined as a softmax function of the difference between the latent prior for each cue with a decision exploration (free) parameter *β*:

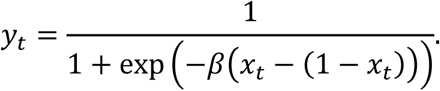

The outcome of a given trial, *o_t_* ∈ {0,1}, is given by *o_t_* = Bernoulli(*σ*(*x_t_*)), where *σ*(*x_t_*) = 1/1 + *e*^−*x*^*^t^*. In our case, we assume that the cue validity of each trial provides evidence for a mutually exclusive state of whether blue is currently the best cue such that:

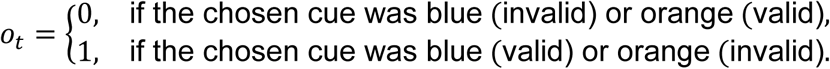

Importantly, priors are estimated based only on past choices {*y_t_*_−1_, *y_t_*_−2_, …} and outcomes {*o_t_*_−1_, *o_t_*_−2_, …}. Therefore, the choice, *y_t_*, and outcome, *x_t_*, of a given trial are only taken into account in the next prior estimate, *x_t_*_+1_.

For the subsequent analysis, we used the prior of the chosen cue being the best, prior = *x_t_* ⋅ I(chosen cue = Blue) + (1 − *x_t_*) ⋅ I(chosen cue = Orange), where I(⋅) equals 1 if the condition inside is true and 0 otherwise. The prior was then categorized into low and high using a subject-level median split.

### Model Fitting and Comparison

To evaluate the performance of the VKF, we consider three alternative learning models: the Rescorla-Wagner (RW; Rescorla and Wagner, 1972), the Kalman Filter (KF; Kalman, 1960), and the Hierarchical Gaussian Filter (HGF; Mathys et al., 2011, 2014). While the VFK, KF, and HGF models account for volatilities in the environment, the RW model does not. A softmax function with a free temperature parameter, *β,* was used as a response model for all learning models as described for the VKF.

The models were fit and compared (**Supplemental Fig. S1**) with a hierarchical Bayesian inference approach based on variational Bayes (Piray et al., 2019). In this approach, fitted parameters are assumed to be drawn from a normal distribution based on the group-level statistics and the model identity is considered a random effect. For all but one subject, the VKF was the best fitting model. Subject-level posteriors were estimated via Laplace approximation. Because participants differ in their learning trajectories, their prior strength distributions vary widely, making fixed or continuous analysis difficult to compare. To assess whether participants integrated priors flexibly according to their relative strength, we compared their behavioral performance between weak and strong priors via a subject-level median-split (see **Supplemental Fig. S2** for individual prior strength trajectories and distributions).

### EEG Data Acquisition and Preprocessing

Electrophysiological (EEG) data were continuously recorded with 257 scalp electrodes (Electrical Geodesics, Inc.) with a 1000 Hz sampling rate and an online reference at Cz. The data were preprocessed using the FieldTrip (Oostenveld et al., 2010) toolbox for Matlab by applying a bandpass Butterworth filter (0.3 to 40 Hz), demeaning, and re-referencing to the common average. Trials with muscle artifacts were visually identified and excluded. Eye and muscle artifacts were manually removed using Independent Component Analysis. The data were down-sampled to 500 Hz, epoched from -1000 to 1000 ms relative to the RDK onset, and z-scored using the trial-wise mean and standard deviation from the -400 to 0 ms interval relative to the cue presentation (before participants could form a specific expectation of the motion direction).

### Univariate Analysis

To test differences in overall channel activity across conditions, time-locked responses were calculated per participant and electrode by averaging across different experimental conditions relative to RDK or response onset. Differences between event-related potentials (ERPs) were assessed using cluster-based permutation statistics as implemented in FieldTrip (Oostenveld et al., 2010).

### Multivariate Analysis

To test whether action and sensory information are shaped by prior strength, we decoded either action (left or right button press) or sensory-like activity (left or right motion direction) from 257 channels for each time point using a Linear Discriminant Analysis (Fisher, 1936; Zhao et al., 2024) via the MVPA-Light (Treder, 2020) toolbox for Matlab. For all decoding analyses, we performed a 10-fold cross-validation procedure. Within each training fold, we balanced the training samples for the selected feature (left/right motion direction or button press), the prior strength (weak or strong), and the cue validity (valid or invalid) using upsampling. The true class probability (TCP) of testing samples was estimated based on the multivariate distribution of the training fold. The final decoding performance was calculated by averaging the TCP values across the 10 testing folds.

To compare the evolution of action and sensory information from cue presentation to motion processing, we performed a time generalization analysis. We trained and tested the generalization of the classifier across all time points for each trial, resulting in a *N_trials_* × *N_training time_* × *N_testing time_* TCP matrix per participant. After training, we excluded trials in which participants incorrectly judged motion direction, as these trials combined expected action with an unexpected sensory outcome or vice versa. This ensured that valid trials reflected expected action and sensory, and invalid trials reflected a joint violation of action and sensory expectations rather than sensory expectation alone. Trials were averaged across conditions, resulting in a *TCP_prior strengt_*_ℎ,*validity*_ matrix with *^N^training time* ^× *N*^*testing time* ^per participant.^

To test the effect of prior strength regardless of its validity (Fig. 3C), we compared the generalization effect of weak priors with strong priors within each validity condition as: Prior Strength Effect = [TCP_Strong, Valid_ − TCP_Weak, Valid_] + [TCP_Strong, Invalid_ − ^TCP^Weak, Invalid^].^

To test whether expected or unexpected action and sensory neural representations are preferentially stabilized, we compared valid (expected) and invalid (unexpected) trials within strong priors (Fig. 4A), Modulation = [TCP_Strong, Valid_ − TCP_Strong, Invalid_]. Preferential stabilization within weak conditions is reported in the

### Supplemental Fig. S4

To test whether validity modulation of action and sensory dynamics under strong priors predict behavior, we correlated the preferential stabilization with behavior across participants. On the neural level, we defined preferential stabilization of expected sensory as the difference between valid and invalid generalization performance (*PG_sensory_* = [TCP_Strong, Valid_ − TCP_Strong, Invalid_]), while we defined preferential stabilization of expected action as (*PG_action_* = [TCP_Strong, Invalid_ − TCP_Strong, Valid_]). On the behavioral level, we reduced participant’s individual differences (Fig. 1C/D), into two dimensions by projecting them into the regression axis between correlated features to capture their shared variance. Cue bias was defined as the coupling between accuracy (Δ*ACC* = *ACC_Strong_*_,*Valid*_ − *ACC_Strong_*_,*Invalid*_) and reaction time modulations (Δ*RT* = *RT_Strong_*_,*Valid*_ − *RT_Strong_*_,*Invalid*_) under strong priors. Precision-weighted prior bias was defined as the relationship between costs in accuracy (*Cost_ACC_* = *ACC_Weak_*_,*Invalid*_ − *ACC_Strong_*_,*Invalid*_) and benefits in RT (*Benefit_RT_* = *RT_Weak_*_,*Valid*_ − *RT_Strong_*_,*Valid*_) when priors were strong relative to weak priors. Neural and behavioral measures were then correlated across participants. This procedure was repeated for response-locked data. Preferential stabilization within weak priors and their behavioral correlations are reported in **Supplemental Fig. S4**.

As the decoding weights cannot be directly interpreted as source activation, the decoded activation patterns were calculated using model inversion as proposed in Haufe et al. (2014). For each time point, activation patterns (A) were reconstructed using the decoding weights (W), *A* = ∑*_x_W*∑^−1^, where ∑*_x_* is the covariance matrix of the neural data, *x*, and ∑^−1^ is the covariance of the estimated latent factors, *Ŝ*, recovered by *Ŝ* = *W^T^x*. Decoding activation patterns were z-scored across space and time for each participant and averaged across participants and time windows of 500 ms between -1.5 and .5 s from RDK onset. For visualization only, electrodes outside the skull area were omitted.

### Statistical Analysis

Unless stated otherwise, we employed paired t-tests. For behavioral data, we tested whether the effects of cue validity on motion accuracy and reaction time depended on prior strength using linear mixed-effects models, with prior strength, cue validity, and their interaction as predictors. Separate models were fitted for accuracy and reaction time, with random intercepts for each participant to account for individual differences. To assess individual differences in behavioral performance, we correlated the effect of cue validity on accuracy (ACC) and reaction time (RT) across participants in the strong prior trials. Cue effects were calculated as: Δ*ACC* = *^ACC^Strong*,*Valid* ^− *ACC*^*Strong*,*Invalid* ^and Δ*RT* = *RT*^*Strong*,*Valid* ^− *RT*^*Strong*,*Invalid*^. For RT, we^ used only correct trials. This effect was quantified specifically for strong prior trials to capture the influence of a reliable predictive cue.

We also explored individual differences in the effect of prior strength by comparing ACC and RT separately for valid and invalid trials. For that, we correlated four metrics: accuracy benefit (*Benefit_ACC_* = *ACC_Strong_*_,*Valid*_ − *ACC_Weak_*_,*Valid*_), accuracy costs (*Cost_ACC_* = *ACC_Weak_*_,*Invalid*_ − *ACC_Strong_*_,*Invalid*_), reaction time benefits (*Benefit_RT_* = *^RT^Weak*,*Valid* ^− *RT*^*Strong*,*Valid*^), and reaction time costs (^*^Cost^RT* ^= *RT*^*Strong*,*Invalid* ^−^ *RT_Weak_*_,*Invalid*_). Thus, larger positive values indicated greater benefits or costs of stronger priors. Because we did not have a priori reason to privilege one specific pairing, we examined all pairwise correlations among these measures while correcting for multiple comparisons using a Sidak correction.

All electrophysiological recordings were assessed using cluster-based permutation tests of paired t-tests and correlations as implemented in FieldTrip (Oostenveld et al., 2010). Clusters were determined across space and time (univariate analyses) or across time (multivariate analyses) by summing t-values within the thresholded cluster. The observed cluster was compared to a surrogate distribution (10.000 permutations) where condition labels were randomly permuted between groups. Effect sizes were quantified using Cohen’s d.

## Results

To assess how predictions modulate sensory and action representations, participants completed a cued motion discrimination task with volatile cue validity (**Fig. 1A**). Rather than using a single cue with fixed validity, as typically used in previous studies, here participants chose one of two cues on each trial, and cue validity varied over time. This design allow us to investigate how trial-by-trial changes in prior strength shape behavior and neural representations while sensory evidence strength remained constant. For optimal behavior, participants were required to continuously update their priors about the cue validity in order to guide cue choice in the next trial, and flexibly weight prior information against sensory evidence. The fact that participants choose which cue to sample provided an additional behavioral measure to improve the estimation of their evolving priors.

### Priors track reported confidence and learning

To estimate participants’ prior strength across trials while controlling for individual differences in learning and decision-making, we fit a volatile Kalman filter (Piray and Daw, 2020; **Fig. 1B**) with a softmax response model to participants’ cue choices (orange, blue) and cue outcomes (valid or invalid). Bayesian model comparison favored the volatile Kalman filter over the alternative models: it had an exceedance probability of 1 and the highest subject-level posterior model probability for 95% of participants (**Supplemental Fig. S1**).

To validate the model-derived priors as estimates of participants’ beliefs about cue validity (purple traces in Fig. 1C), we tested whether stronger priors were associated with greater self-reported confidence, the task’s validity structure, and subsequent cue choices. Priors were stronger when participants reported high confidence in the chosen cue (**Fig. 1D**, left; p < .001, t(19) = 8.22, d = 1.84) and during low uncertainty blocks (**Fig. 1D**, center; p < .001, t(19) = 12.30, d = 2.75), which included a cue that indicated the correct subsequent motion direction 80% of the times. During low uncertainty blocks, stronger priors also predicted more frequent selection of the currently optimal cue (**Fig. 1D**, right; median split; p < .001, t(19) = 14.71, d = 3.29). These findings support the model-derived priors as a measure of participants’ latent belief about the cue validity across contingency reversals.

### Prior strength increases the influence of cue information during motion decisions

Having established that the model estimates tracked participants’ cue beliefs, we next asked whether these priors influenced the subsequent decision. We tested whether priors strength modulated the effect of the cue validity on motion accuracy (ACC) and reaction time (RT) using trial-level mixed effect models. To define weak and strong priors relative to each participant’s own prior range, we use a within-participant median split of prior strength. Stronger priors elicited stronger cue validity effects for both accuracy (**Fig. 2A**; p < .001, t(6004) = 10.19; f^2^ = .019) and reaction time (**Fig. 2B**; p < .001; t(6004) = 7.19; f^2^ = .011).

**Figure 2.**
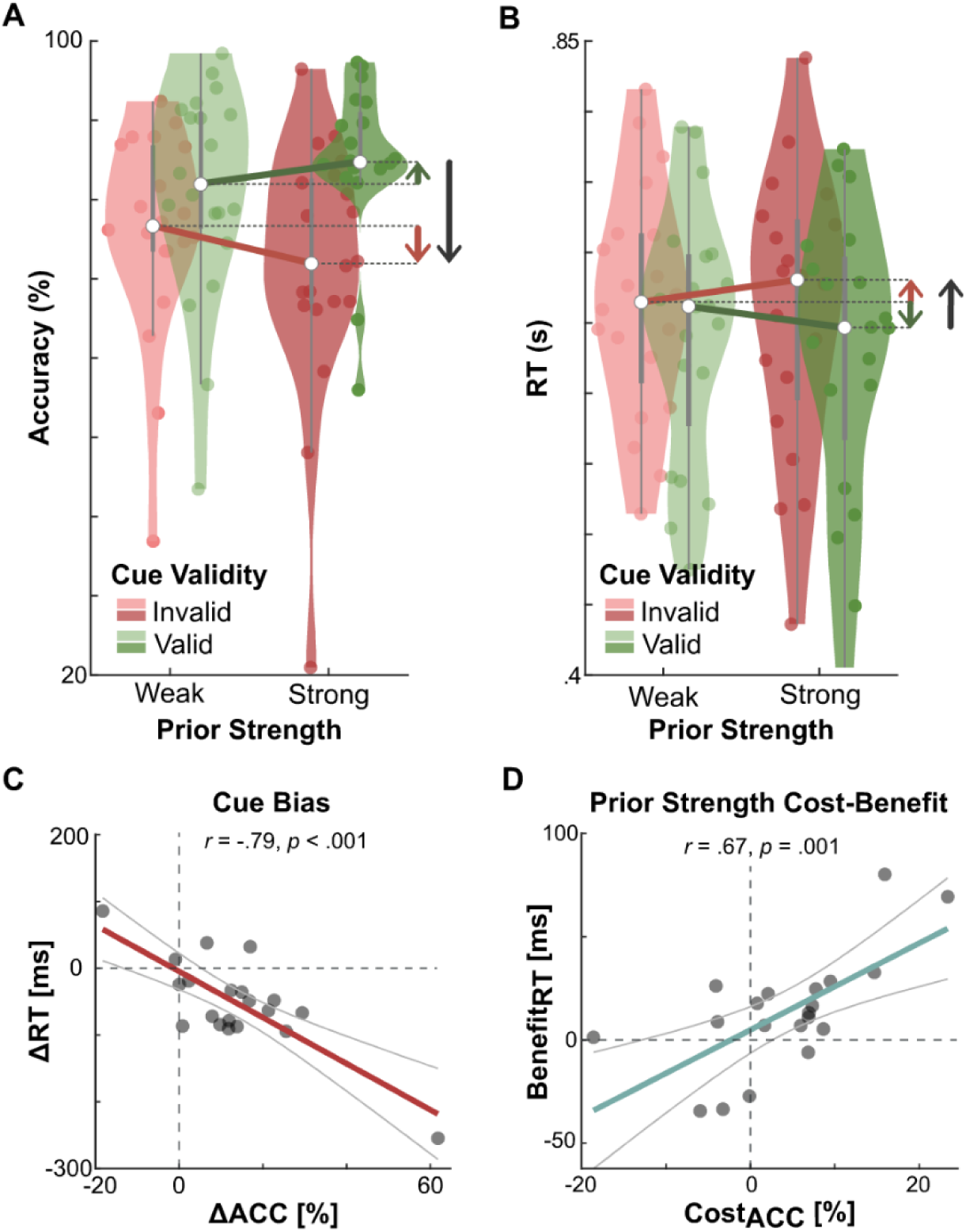
Priors induced cue bias and precision-weighted prior bias. (**A**) Accuracy (ACC) of motion choices as a function of cue validity and prior strength. Green and red arrows indicate group-level benefit (*Benefit_ACC_*) and cost (*Cost_ACC_*) in accuracy of strong prior trials within the same cue validity. Black arrows indicate the modulation in accuracy by cue validity on strong prior trials (*ΔACC* = *ACC_Strong_*_,*Valid*_ − *ACC_Strong_*_,*Invalid*_). (**B**) Same conventions as in panel (A) for reaction time (RT) instead of accuracy. (**C**) Correlation between *ΔACC* and *ΔRT* (correct trials only) across subjects. (**D**) Correlation between RT benefits (*Benefit_RT_* = *RT_Weak_*_,*Valid*_ − *RT_Strong_*_,*Valid*_) and accuracy costs (*Cost_ACC_* = *ACC_Weak_*_,*Invalid*_ − *ACC_Strong_*_,*Invalid*_). Dots represent individual participants.

We further examined the effect of prior strength separately for fulfilled (valid) and violated (invalid) predictions. Stronger priors resulted in a behavioral benefit on valid trials, increasing accuracy by 4% (SD = 6.6%) and reducing reaction times by 16 ms (SD = 27 ms) relative to weak priors. These effects were significant at the group level for both accuracy (p = .011, t(19) = 2.82, d = .34), and reaction time (p = .017, t(19) = −2.62, d = .17) In contrast, when predictions were violated, stronger priors were associated with a 4% decrease in accuracy (SD = 9%) and a small, inconsistent 7-ms change in reaction time (SD = 29 ms) relative to weak priors. Neither effect during invalid trials reached statistical significance (accuracy: p = .056, t(19) = −2.04, d = .28; reaction time: p = .271, t(19) = −1.13, d = .08). Together, these results suggest that the influence of prior strength on behavior was more consistent across participants when predictions were fulfilled than when they were violated.

### Individual differences in behavior reflect cue bias and precision-weight prior bias

Having established that prior strength modulated behavior at the group level, we next examined individual differences in these effects. We first quantified cue bias under strong priors to best capture the impact of a reliable cue. Cue bias was indexed by the across-participants correlation between changes in accuracy (Δ*ACC* = *ACC_Strong_*_,*Valid*_ − *ACC_Strong_*_,*Invalid*_) with changes in RT (Δ*RT* = *RT_Strong_*_,*Valid*_ − *RT_Strong_*_,*Invalid*_; correct trials only). A statistically significant negative correlation (**Fig. 2C**; r = -.79, p < .001) indicated that some participants relied more than others on the cue information. We denoted the shared variance between these measures as ***cue bias***, reflecting how strongly behavior was biased toward strongly predictive cues.

Following on individual differences in cue bias, we explored individual differences in how strongly their behavior changed with prior strength. For each participant, we estimated accuracy benefits (*Benefit_ACC_* = *ACC_Strong_*_,*Valid*_ − *ACC_Weak_*_,*Valid*_), accuracy costs (*Cost_ACC_* = *ACC_Weak_*_,*Invalid*_ − *ACC_Strong_*_,*Invalid*_), reaction time benefits (*Benefit_RT_* = *^RT^Weak*,*Valid* ^− *RT*^*Strong*,*Valid*^), and reaction time costs (^*^Cost^RT* ^= *RT*^*Strong*,*Invalid* ^−^ *RT_Weak_*_,*Invalid*_), and correlated these four metrics across participants. The only association that survived correction for multiple comparisons was a positive correlation between accuracy costs and reaction time benefits (**Fig. 2D**; r = .67, p < .001). Participants who gained greater speed benefits from strong valid priors also made more errors when those priors were invalid. We denoted the variance along this shared dimension as ***precision-weighted prior bias***, reflecting the extent to which participants scaled prior reliance with prior strength. This is an important distinction for Bayesian theories, which predict a reliability-weighted prior reliance.

### Priors distinctively stabilize sensory and action information

To investigate how priors shape neural dynamics, we used uni- and multivariate analyses of hd-EEG. Event-related potentials locked to the RDK onset revealed no significant effects of prior strength in either validity condition (**Supplemental Fig. S3**), suggesting that stronger learned priors did not produce changes in mean evoked activity in specific channel clusters. However, prior strength may shape distributed neural representations and not solely the mean activity (de Lange et al., 2018; Kok et al., 2012; Nigam & Schwiedrzik, 2024; Tang et al., 2018a). Hence, we implemented a time-resolved multivariate decoding approach to test whether prior strength shaped sensory (motion direction) and action information (button response) in using a linear discriminant analysis (Fisher, 1936) across all channels. Decoding was implemented using a 10-fold cross-validation procedure controlling for cue validity and prior strength. After training, condition comparisons were restricted to trials with correct motion responses, allowing valid and invalid trials to be interpreted as expected and unexpected action/sensory information, respectively.

Sensory and action information were reliably decoded after stimulus onset (**Fig. 3A/B**; *p_sensory_* = .041, *t̄_cluster_* = 2.77, d = .78; *p_action_* = .001, *t̄_cluster_* = 3.48, d = 1.25), and around the response (*p_sensory_* = .004, *t̄_cluster_* = 3.40, d = .95; *p_action_* = .001, *t̄_cluster_* = 4.59, d = 1.62). Action information was also decodable during early stimulus presentation (–20 to 86 ms relative to RDK onset, p = .047, *t̄_cluster_* = 2.44, d = .63), suggesting anticipatory action information processing. Sensory activation patterns (Haufe et al., 2014) were more pronounced over posterior regions (**Fig. 3C**), while activation patterns for action information were stronger over fronto-central sensors (**Fig. 3D**), in line with the expected cortical organization of visual and motor cortices.

**Figure 3.**
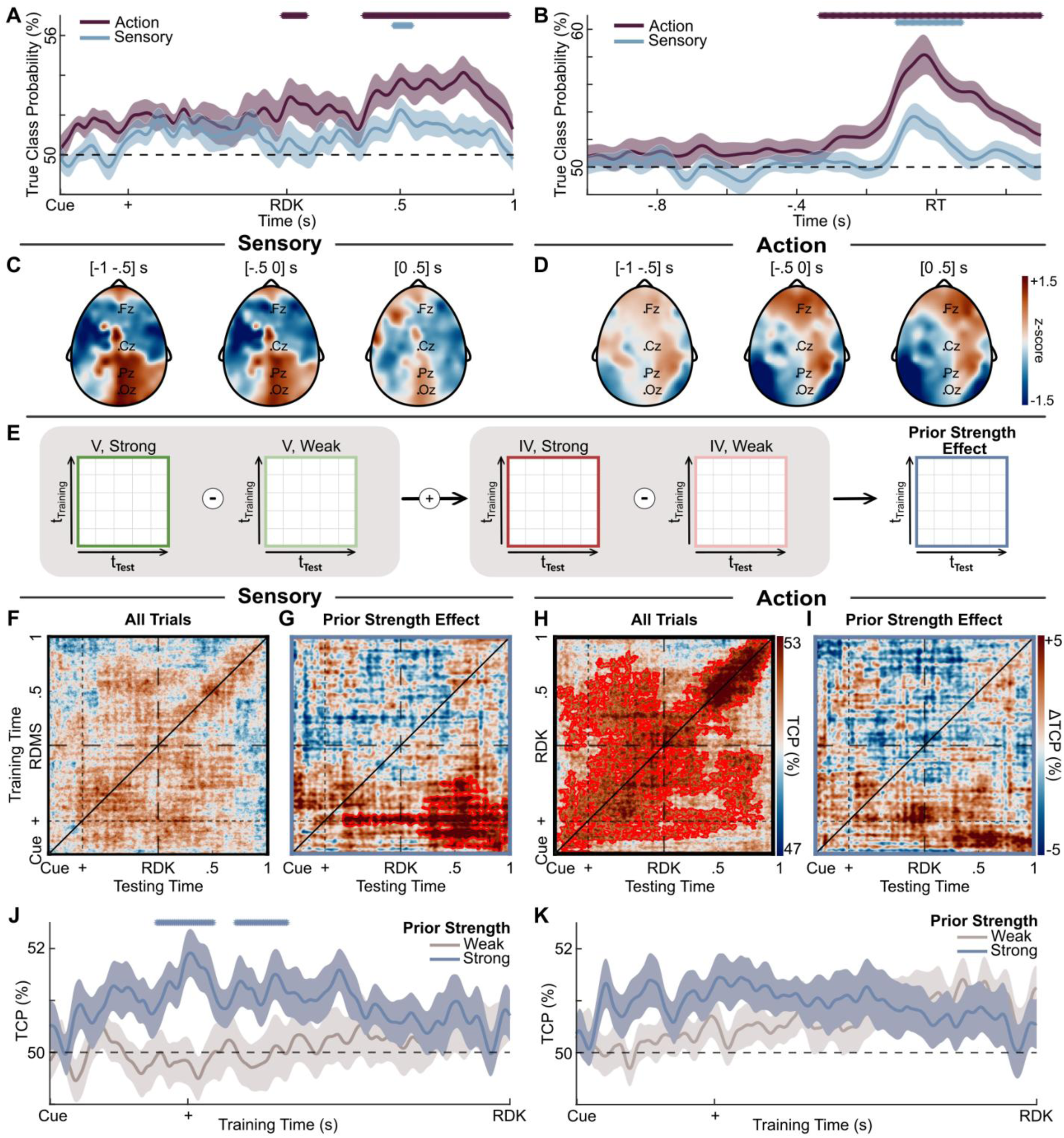
Stronger priors enhance the generalization of pre-stimulus sensory activity. (**A**) Average true class probability (TCP) of action (left/right response) or sensory (left/right motion) information across subjects, aligned to stimulus onset. Data was smoothed with a Gaussian-weighted average over a window of 100 ms for visualization. Shaded areas show ±1 SD. (**B**) Same data as (A), but time-locked to the reaction time. (**C**) Activation patterns for decoded action and (**D**) sensory information. (**E**) Isolating the effect of prior strength from validity: computed as the sum of differences in TCP between strong and weak prior trials for both valid (V) and invalid (IV) conditions, averaged across subjects. (**F**) Average cross-decoding performance of sensory information across all conditions. (**G**) Estimated effect of prior strength across time for sensory information. (**H-I**) same as (F-G) but for action information. (**J**), Average TCP across all testing times for pre-stimulus training times comparing weak and strong prior trials for sensory and (**K**) action decoding. Top lines indicate the time points of significant clusters. Red contours mark significant positive clusters.

If strong priors increase sensory or action representations, decoding performance should differ between weak- and strong-prior trials before stimulus onset. We tested this separately for valid and invalid trials as validity determines whether the sensory evidence and corresponding action confirms or violates the expectation. Collapsing across validity conditions could therefore obscure different effects of priors on expected and unexpected representations. Contrary to this hypothesis, time-resolved decoding did not statistically differ between strong- and weak-prior trials at any pre-stimulus time point, for either valid or invalid trials, in either sensory or action domain (**Supplemental Fig. S4**). Thus, stronger priors did not simply increase the instantaneous expression of sensory or action information.

The absence of instantaneous decoding differences between weak and strong priors does not rule out differences in the temporal evolution of neural representations. For example, stronger priors might lead to more persistent or stable representations, even if they do not lead to stronger instantaneous representations. To test this possibility, we used cross-temporal generalization to examine how sensory and action information evolved over time under strong and weak priors. We first asked whether sensory and action information showed any temporal generalization regardless of prior strength and validity. Across all trials, action information showed a mixture of dynamic and stable coding (King & Daheane, 2014), with strongest decoding along the temporal diagonal and, at the same time, cross-temporal generalization in the last 500 ms of the RDK display (**Fig. 3H**; p = .005, *t̄_cluster_* = 1.36; d = .30). In contrast, sensory information (**Fig. 3F**) showed no reliable cross-temporal generalization across all trials.

Having established domain-specific coding principles across all trials, we asked whether strong priors increase the stability of sensory or action neural information relative to weak priors (**Fig. 3E**). If strong priors stabilize representations regardless of whether incoming sensory evidence confirms or violates the prediction, there should be stronger or broader temporal generalization for strong than weak priors after controlling for the validity effect. Stronger priors enhanced the generalization of pre-stimulus sensory information (p = .043, *t̄_cluster_* = 1.60; d = .36, **Fig. 3G**), while no corresponding effect was observed for action information (**Fig. 3I**). Focusing on the pre-stimulus period, sensory information at the end of the cue presentation (p = .03, *t̄_cluster_* = -3.75; d = .96) and during the delay period (p = .048, *t̄_cluster_* = -2.53; d = .59) generalized more across the trial when priors were strong (**Fig. 3J**), but action information did not (**Fig. 3K**). This suggests that, when priors are strong, information about the cue-driven expected sensory input is preserved in the neural activity across the trial.

Together, these results suggest that the strength of learned priors did not simply increase sensory or action information representations. Instead, strong priors selectively increase the stability of pre-stimulus sensory information over time. Unlike sensory, action information was temporally stable across all trials and not modulated by prior strength.

### Preferential stabilization of neural representations predicts cue bias and precision-weighted prior bias

The previous analyses showed the stability of sensory and action information across prior strength, but it did not indicate whether this stabilization favored expected or unexpected information. This distinction is central to predictive processing theories: Bayesian accounts predict facilitation of expected information processing whereas cancellation accounts predict facilitation of unexpected information processing. To test these predictions, we compared the cross-temporal generalization between valid (expected) or invalid (unexpected) trials under strong priors and tested whether preferential stabilizations were related to behavior (**Fig. 4A**). Greater generalization for valid than invalid trials would indicate a more stable representation of expected information, consistent with Bayesian accounts. In contrast, greater generalization for invalid than invalid trials would indicate a more stable representation of unexpected information, consistent with cancellation accounts.

**Figure 4.**
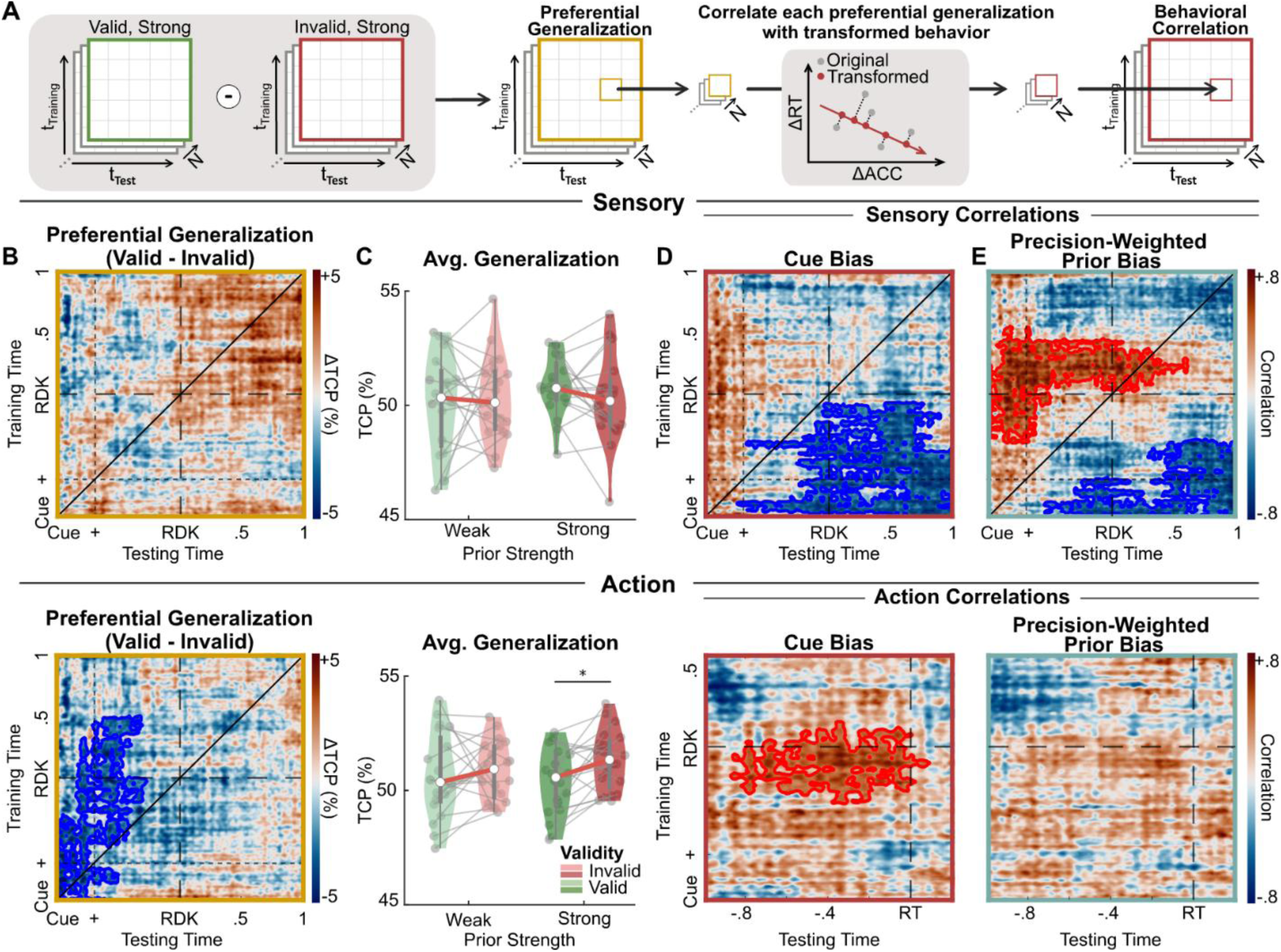
Behavioral correlates of cue bias and precision-weight prior bias. (**A**) Analysis pipeline correlating behavioral performance with preferential generalization during strong prior trials across subjects. (**B**) Grand-mean difference of preferential generalization (*TCP_Strong_*_,*Valid*_ − *TCP_Strong_*_,*Invalid*_) for sensory (top) and action (bottom). Positive values indicate stronger generalization for expected information, while negative values indicate stronger generalization for unexpected information. (**C**) Grand-mean TCP over all training and testing times for each condition. Dots represent individual participants. (**D**) Correlation between cross-temporal sensory (top) and action (bottom) preferential generalization of expected sensory and unexpected action with cue bias and (**E**) precision-weighted prior bias. Red and blue contours mark significant positive and negative clusters, respectively.

Across participants, sensory information exhibited no consistent preferential stabilization under strong priors (**Fig. 4B**, top). Unlike sensory, unexpected action showed greater generalization than expected action information (**Fig. 4B**, bottom; p = .033, *t̄_cluster_* = -1.36; d = .30). This effect was also present when generalization was averaged across all training and testing times (**Fig. 4C**, bottom, p = .008, t(19) = 2.98, d = .667). Importantly, participants could not know that the unexpected action would be the correct response before the stimulus onset. Thus, the effect does not reflect pre-stimulus representations of the future correct action. Instead, it suggests a stable, but possibly decaying, representation of the alternative response that was later recruited when sensory evidence violated the cue. Together, these results address our second hypothesis: strong priors referentially stabilized unexpected action representations, consistent with cancellation accounts, while sensory representations showed no consistent group-level preference.

We next asked whether preferential stabilization predicted behavior across participants, even when no consistent group-level modulation was observed for sensory information. To test this, we correlated preferential stabilization driven by strong priors with cue bias and precision-weighted prior bias across participants. We defined preferential stabilization separately for action and sensory information, based on the direction predicted by the group-level and theoretical effects. For action, we quantified preferential stabilization of unexpected action information (*PG_action_* = *TCP_Strong_*_,*Invalid*_ − *TCP_Strong_*_,*Valid*_). For sensory, we quantified preferential stabilization of expected sensory ^information (^*^PG^sensory* ^= *TCP*^*Strong*,*Valid* ^− *TCP*^*Strong*,*Invalid*^).^

Cue bias was negatively correlated with pre-stimulus *PG_sensory_* (p = .014, *r̄_cluster_* = -.48; **Fig. 4D**, top). Participants with preferential stabilization of expected pre-stimulus sensory information showed reduced cue bias, contrary to Bayesian or OPT predictions. Cue bias was also positively correlated with pre-stimulus *PG_action_* (p = .041, *r̄_cluster_* = .44; **Fig. 4D**, bottom). Specifically, participants with stronger stabilization of unexpected actions before the response showed stronger cue bias. Thus, stronger cue bias was associated with a relative shift toward stabilizing unexpected information in both sensory and action representations, in line with cancellation accounts.

Unlike cue bias, precision-weighted prior bias dissociated across domains. It was not correlated with preferential stabilization of action information. In the sensory domain, the correlation depended on the direction of temporal generalization. Stronger forward *PG_sensory_* predicted weaker precision-weighted prior bias (p = .043, *r̄_cluster_* = -.43). Thus, maintaining the initial expected representation into the stimulus period predicted weaker reliability-dependent prior reliance across participants, contrary to predicted by Bayesian or OPT accounts. The opposite relationship emerged for backward generalization: stronger stimulus-driven *PG_sensory_* predicted stronger precision-weighted prior bias (p = .023, *r̄_cluster_* = .49). Thus, participants whose pre-stimulus activity resembled the sensory representations expressed during subsequent stimulus processing also showed stronger reliability-dependent prior bias, consistent with Bayesian reliability weighting. Such asymmetry in cross-temporal decoding can occur with ramping representations due to differences in signal-to-noise ratio between training and testing times (King & Dehaene, 2014). Since both action and sensory representations are stronger during late RDK (**Fig. 3B**), asymmetric effects are to be expected.

Under weak priors, no corresponding preferential stabilization or behavioral correlations were found (**Supplemental Fig. S4**). Together, these results suggest that strong priors engaged domain-specific predictive mechanisms: cancellation-like action dynamics were associated with cue bias, whereas sensory dynamics showed temporally opposing relationships with precision-weighted prior bias that were compatible with both Bayesian and cancellation accounts.

## Discussion

This study tested whether learned predictions preferentially stabilize expected or unexpected information, and whether these effects differ across sensory and action representations as decision unfold. Behaviorally, participants tracked changes in cue reliability (**Fig. 1D**) and relied more on the cue information when their priors were strong (**Fig. 2A/B**), consistent with Bayesian accounts of decision-making. At the neural level, the predictive effects on temporal stability differed across domains. Sensory and action signals showed different patterns of stability over time, indicating that no single Bayesian or cancellation account can explain predictive processing across both domains.

### Predictive processing for action control

An adaptive action system should use prior information to prepare the most likely response while remaining flexible enough to switch when those expectations are violated. Our results suggest that action representations may strike this balance by combining dynamic and stable coding (**Fig. 3H**), consistent with accounts in which multiple candidate actions are prepared in parallel and continuously updated until one is selected (Cisek, 2007; Gallivan et al., 2018). In this view, stable patterns may keep alternative responses available, whereas changing patterns may reflect ongoing adjustments as new evidence comes in.

Under strong priors, we observed that representations of the unexpected action were more stable than those of the expected action (**Fig. 4B**, bottom). Critically, the size of this effect predicted participants’ cue bias (**Fig. 4D**, bottom). The participants who relied more on the cue also showed greater stabilization of the alternative, unexpected action. This pattern is consistent with cancellation accounts, in which predictions can enhance processing of deviations from expectation. Importantly, preferential generalization of unexpected actions was associated with cue bias but not with precision-weighted prior bias (**Fig. 4E**, bottom). This observation suggests that action representations reflect stronger predictive information irrespective of individual differences in how strongly prior reliance scales with prior reliability. Previous findings had linked prior expectations to motor preparation (e.g., de Lange et al., 2013; Kelly et al., 2021; Michaels et al., 2025). Our findings extend this view by suggesting that predictive processing also helps keep an alternative response “ready” when expectations are violated, supporting rapid execution of alternative actions.

### Predictive processing for sensory inference

An adaptive sensory system faces a similar challenge: it should use prior information to narrow down likely interpretations while remaining sensitive to new evidence that may contradict those expectations. Our results show that sensory representations mainly rely on a dynamic code, yet strong priors stabilized pre-stimulus sensory information (**Fig. 3G**). This finding is consistent with studies showing that expectations decreased local neural variability (Geurts et al., 2022; Skowron et al., 2024) and inter-areal correlations (van Bergen and Jehee, 2018; Lange and Haefner, 2022; van den Brink et al., 2023).

Although this increase in stability by strong priors did not consistently favor expected or unexpected information (**Fig. 4B**), its temporal evolution revealed two opposing relationships with prior reliance (**Fig 4E**, top). On one hand, pre-stimulus activity in participants with stronger precision-weighted prior bias already resembled the subsequent expected sensory representation (backward generalization). This finding is consistent with Bayesian accounts and can be conceptualized as pre-stimulus sensory templates (Kok et al., 2014, 2017).

On the other hand, stronger stability of cue-driven expected sensory representation during stimulus processing predicted weaker cue bias and precision-weighted prior bias (forward generalization). Notably, this finding is inconsistent with a Bayesian account in which enhanced processing of expected information leads to greater reliance on the prior.

One possibility is that this negative relationship reflects a cancellation process in the sensory domain, in which stronger generalization of unexpected sensory information predicts stronger prior reliance. Alternatively, it may be understood within a predictive coding account (Furutachi & Hofer, 2026; Rao & Ballard, 1999), in which predictions provide a top-down reference template to which sensory input can be compared. Under this interpretation, a more stable template might facilitate mismatch detection, thereby supporting the correction of invalid priors. This interpretation is compatible with, but not directly demonstrated by, observations of high-level prediction errors (Heilbron & Lange, 2025; Richter et al., 2024) and stimulus-specific sensory amplification of unexpected information (Furutachi et al., 2024).

### Convergences and divergences between sensory and action predictive processing

Predictive signals must coordinate multiple processing stages: from interpreting incoming sensory evidence to selecting and implementing an action. Sensory and action representations were shaped similarly by learned predictions in two ways: a) priors shaped the stability of representations over time more consistently than their representation strength and; b) cue bias was correlated with generalization of pre-stimulus unexpected information, congruent with cancellation accounts. These convergences suggest that both sensory and action systems remain sensitive to alternative choices, in case the prior is violated. This mechanism might explain how predictions lead to stronger responses to unexpected stimulus (Furutachi et al., 2024; Tang et al., 2018a, 2023) and preparation of alternative actions (Gallivan et al., 2018).

Sensory and action representations also exhibited several differences. First, action representations were stable across the trial regardless of prior strength, whereas sensory representations were stabilized only when priors were strong. Second, participants showed a tendency to preferentially stabilize unexpected actions. Third, only sensory representations predicted precision-weighted prior bias across participants. Together, these observations suggest that sensory representations were more sensitive to the reliability of the prior, consistent with precision-weighted accounts of perceptual inference (e.g. Feldman & Friston, 2010).

### Limitations and Future Directions

Previous work had quantified neural representations of expected or unexpected information (de Lange et al., 2018; Press et al., 2020) as event-related responses (Alink et al., 2010; Richter et al., 2018), tuning curves (Park et al., 2023), representational strength (Kok et al., 2012, 2014; Richter et al., 2022) or the stability of expected information (Kok et al., 2017; Tang et al., 2018b). Most studies focus on visual processing, but prior representations are widespread across the neocortex (Findling et al., 2025). Our findings extend this work by showing that predictive processing differs across sensory and action representations.

At the same time, our design cannot fully address several additional questions. First, participants used a fixed stimulus-response mapping, raising the question of whether action dynamics reflected a specific motor plan, an abstract choice, or the transformation between the two. Second, participants selected which cue to sample, introducing a exploration-exploitation trade-off (Domenech et al., 2020) that might have shaped how priors were used. Finally, discretizing priors into weak and strong may have obscured nonlinear relationships (Foucault et al., 2024).

Beyond these design choices, the volatility of the task may have increased individual variability in predictive processing. Unlike studies with more stable contingencies, participants had to continuously update the reliability of their predictions, rather than maintain a fixed expectation. This may explain both the absence consistent differences in evoked responses and the lack of consistent preferential stabilization of sensory information across participants. Such variability may arise from differences in learning strategies under uncertainty, the timescale over which priors impact perceptual decisions (Gold and Stocker, 2017), or how priors are embedded in the transformation of sensory evidence (Sohn et al., 2019).

The lack of consistent group-level effects raises a broader question of whether short-term priors rely on mechanisms distinct from long-term priors. Consistent with this hypothesis, stronger responses to expected sensory information has been reported only after extended task exposure (Walsh et al., 2024), and expectation suppression effects were observed following prolonged training (Feuerriegel et al., 2021). A testable prediction is that long-term priors should produce more consistent group-level stabilization of expected sensory information, whereas rapidly changing priors should preserve greater individual variability and stronger dependence on trial-by-trial reliability. Such comparisons can clarify how short-term predictive inference relates to long-term statistical learning thought to optimize sensory coding in natural environments (Simoncelli and Olshausen, 2001; de Lange et al., 2018).

## Conclusion

Our results show that predictive processing is not captured by a single Bayesian or cancellation model. Action representations primarily followed a cancellation-like pattern that stabilized unexpected information, whereas sensory representations showed stage-dependent dynamics consistent with both Bayesian and cancellation accounts. These findings suggest that these mechanisms are complementary components whose expression depends on domain and decision stage.

## Author contributions

Conceptualization: GI, RFH; Methodology: GI, RFH; Formal Analysis: GI; Investigation: GI; Resources: RFH; Writing – Original Draft, GI; Writing – Review & Editing: GI, AB, RFH; Supervision: AB, RFH; Funding Acquisition: GI, RFH.

## Conflict of interest

The authors declare no competing interests.

## Acknowledgments

This work was funded by the doctoral scholarship from the Baden-Württemberg Landesgraduiertenförderungsgesetz (GI), the International Max Planck Research School for The Mechanisms of Mental Function and Dysfunction (GI), the German Research Foundation (HE8329/2-1 to RFH), the Medical Faculty of the University of Tübingen (JRG Plus program, RFH), and the Jung Foundation for Research and Science (Ernst Jung Career Advancement Award in Medicine; RFH). The funders had no role in the study design, data collection, analysis, decision to publish, or preparation of the manuscript.

## Notes

### Competing Interest Statement

The authors have declared no competing interest.

### Summary of Updates

Manuscript was updated to improve clarity. Authors were updated.

