## Supplemental Figures for "Predictive processing for sensory and action neural representations"

**Supplemental Figure 1**

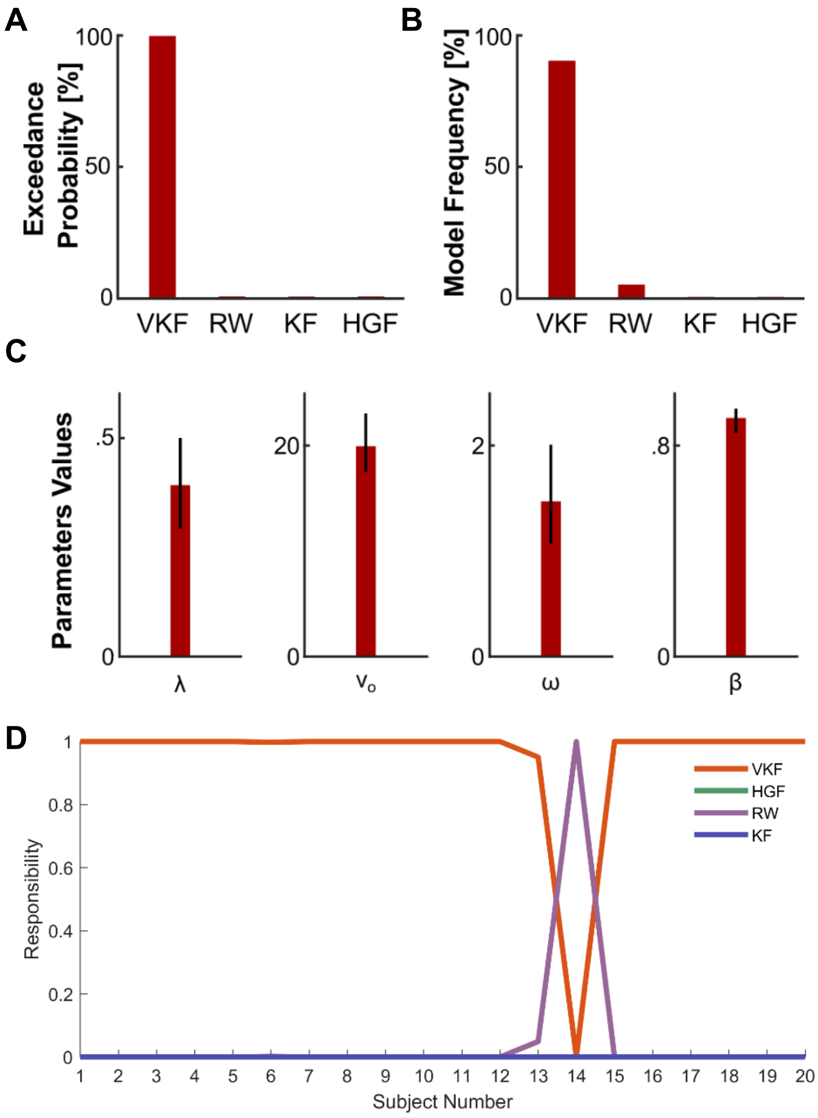


**Figure S1. Model comparison between the Volatile Kalman Filter (VKF), Rescorla-Wagner (RW), Kalman Filter (KF), and Hierarchical Gaussian Filter (HGF) models.** (**A**), Exceedance probability, the probability that a model is the most likely between the alternative models. (**B**), Frequency in which a model was the best-fitting one between the alternative models. (**C**) Average and standard deviation of parameter values for the VKF. (**D**) Model responsibility across subjects, the probability that a model explains each subject's data given all models tested.

### Supplemental Figure 2

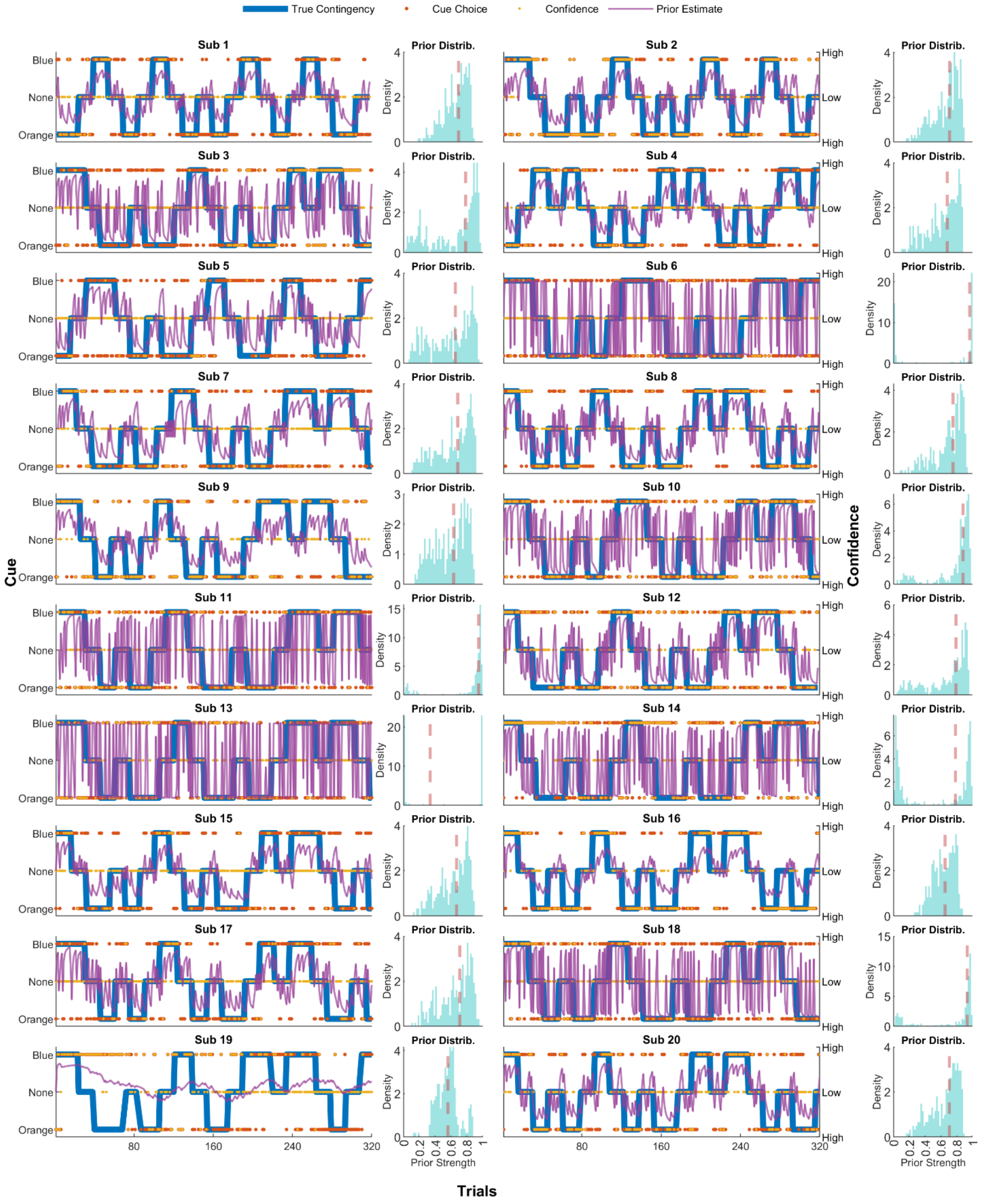


**Figure S2. Single subject behavioral data (left) and distribution of prior strengths for chosen cues across trials (right).** Best cue (true contingency; blue), cue choice ($y_t$, orange), outcome ($o_t$; yellow), and estimated prior from VFK ($x_t$; purple) across trials. Red dashed line indicates the median split.

### Supplemental Figure 3

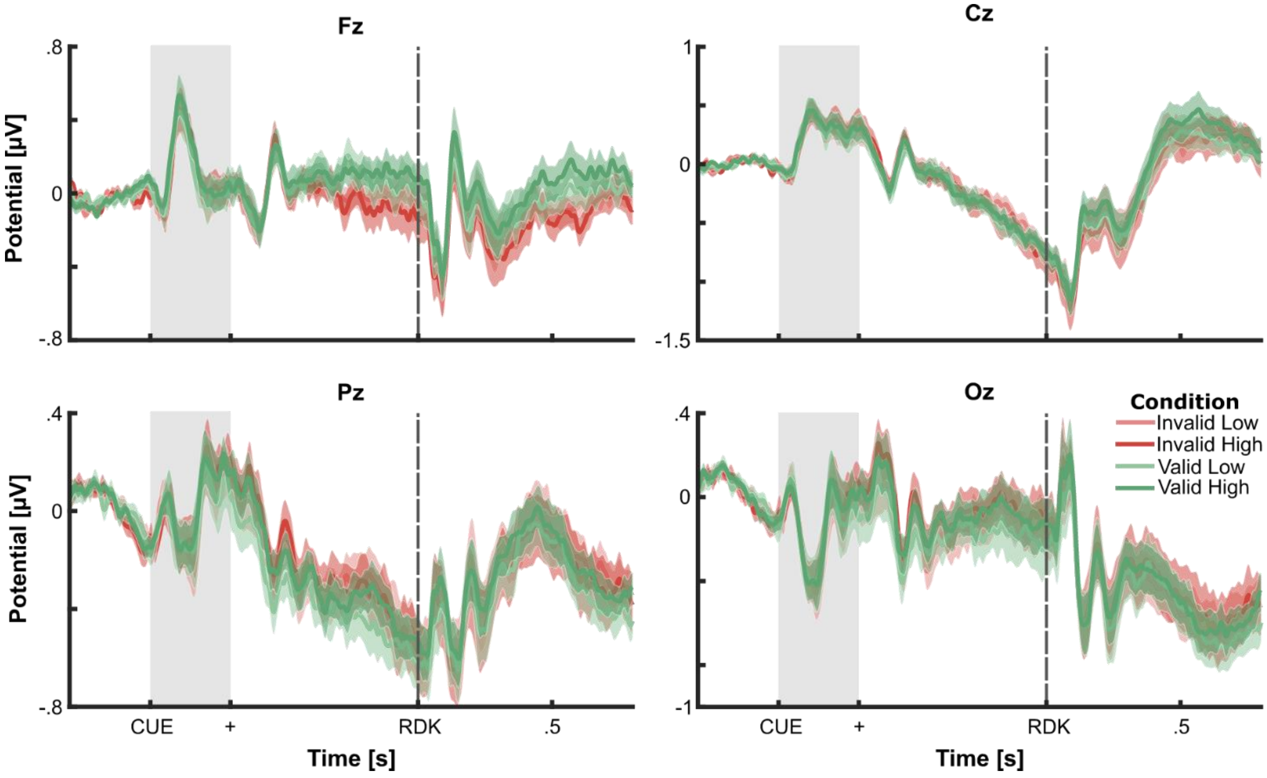


**Figure S3. Event-related potential of central electrodes (Fz, Cz, Pz, and Oz) of belief (high, low) and validity (valid, invalid) conditions.** The data was averaged across participants and time-locked to the Random Dot Kinematogram (RDK) onset and baseline corrected to pre-cue onset activity. Shaded areas show the standard deviation from the mean.

### Supplemental Figure 4

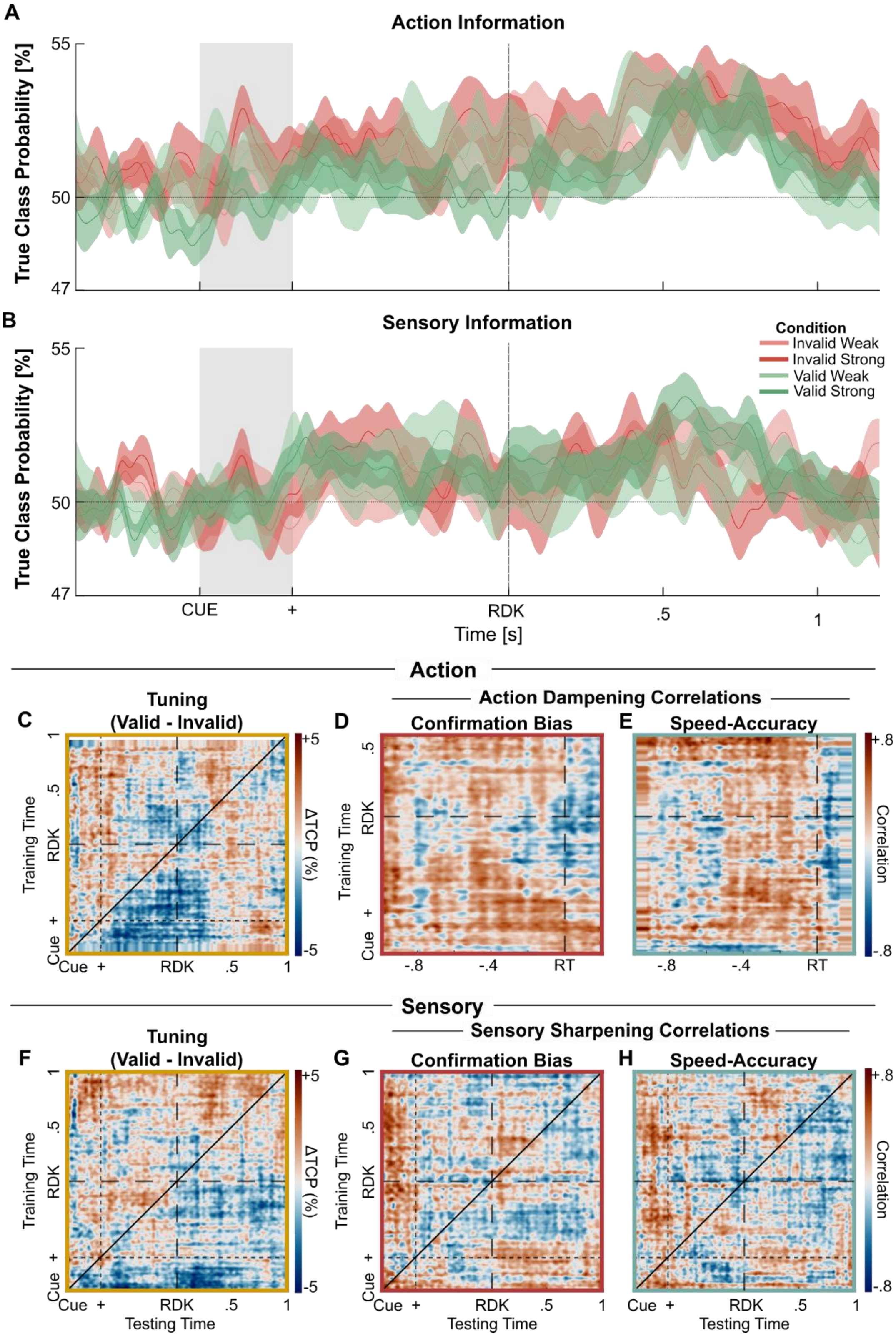


**Figure S4. Decoding performance across conditions and temporal generalization for weak priors.** (**A**) True class probability across conditions across prior strength (high, low) and validity (valid, invalid) conditions for action and (**B**) sensory information. The data was averaged across participants and time-locked to the Random Dot Kinematogram (RDK) onset. Shaded areas show the standard deviation from the mean. (**C**) Grand-mean difference between the cross-temporal generalization of valid and invalid low prior trials ($TCP_{Low,Valid} - TCP_{Low,Invalid}$), reflecting tuning by prior validity for action information. (**D**) Correlation between cross-temporal action dampening with confirmation bias and (**E**) with speed-accuracy trade-off. (**F–H**) Same conventions as for panels (C–E), but for sensory information.
